# Affordable plasmonic biosensing: democratizing SERS with scalable, field-compatible substrate fabrication

**DOI:** 10.1101/2025.06.05.658202

**Authors:** Saransh Arora, Vighnesh Ginde, Swati Tanwar, Marwan El Chazli, Raj Bhatt, Evan Edelman, Debadrita Paria, Ishan Barman

## Abstract

Efficient and accurate plasmonic biosensing in the field remains a significant challenge. Despite its potential to revolutionize point-of-care (PoC) diagnostics through its unparalleled sensitivity and precise molecular fingerprinting capabilities, adoption of traditional surface-enhanced Raman spectroscopy (SERS) in the field has not proven feasible yet. High production and material costs, complex fabrication methods, reliance on specialized equipment, and persistent issues with sensor stability and reproducibility continue to impede development for PoC use. Addressing these challenges, this study innovates a democratized and cost-effective fabrication kit that enables the production of SERS substrates using commonly available materials and straightforward electrochemistry techniques without compromising on sensitivity and reproducibility. Importantly, this method leverages commercially available bottled water and simple battery-powered fabrication, thereby eliminating reliance on a power grid and enabling the local production of biosensors in resource-restricted and conflict-affected areas. The cost of producing the fabrication kit is $39.54 with raw materials purchased at bulk retail prices, while the cost of consumables for fabricating each test is just 1.33¢. To ensure real-world feasibility, we conducted a comprehensive reproducibility analysis, where consistent plasmonic enhancement was observed across multiple production batches. Furthermore, we demonstrated their efficacy in two critical applications: the rapid detection of bacteria and pesticides. We detect trace levels of pesticides such as Thiram and Thiabendazole down to 0.1 ppm using a digital SERS approach. We also demonstrated the identification of bacteria isolated from culture, namely *Escherichia coli* and *Bacillus subtilis*. We envision that this label-free, high-sensitivity substrate, when paired with portable Raman spectrometers, could open doors for a new era of field-deployable biosensing, paving the way for its adoption for a plethora of applications, from public health to food testing.

## 1. Introduction

The COVID-19 pandemic has underscored the urgent need for innovative testing platforms that can advance global health equity. In a time of crisis, while high-income nations raced to develop solutions and stockpile critical resources, low-income regions and displaced communities, particularly those affected by armed conflict, were often left without the means to respond effectively. The world needs low-cost, high-sensitivity biosensors that can be manufactured from readily available materials and used in point-of-care settings. Many of these fragile areas lack functional hospitals, testing facilities, trained medical personnel, and reliable WASH (water, sanitation, and hygiene) services, forcing them to rely on makeshift facilities. In addition, frequent power outages jeopardize temperature-sensitive reagents, poor road networks make timely sample transportation difficult, and a lack of water purification systems make testing production in these regions next to impossible^1–3^.

Among emerging solutions, surface-enhanced Raman spectroscopy (SERS), the plasmonic counterpart of conventional Raman spectroscopy, shows exceptional promise for creating robust, low-cost, and high-sensitivity testing platforms. By leveraging near-field enhancements from plasmonic nanostructures to amplify weak Raman signals, SERS offers unparalleled sensitivity, specificity, non-bleachable detection, and a rapid time-to-result, making it particularly well-suited for resource-limited settings where rapid, adaptable testing is crucial^4–7^. Unlike conventional techniques, label-free SERS does not require additional reagents, allowing testing kits to be rapidly adapted to new analytes or challenges with minimal modifications, such as a software update. Despite its immense potential and growing popularity in preclinical research, the widespread adoption of SERS in real-world applications has been hindered by high production costs, complex substrate fabrication processes, and specialized training needs.

Realizing the full potential of SERS-based mass screening in the field hinges on several critical design criteria. First, the sensor must provide high optical enhancement to achieve a low detection limit. Second, it should support label-free detection and broad applicability, minimizing the need for analyte-specific reagents. Third, multiplexing capabilities are essential, along with resilience against interference from contaminants and variations in handling, storage, and transportation. Fourth, the cost of production setup, raw materials, and fabrication of each test must be low to make the test affordable globally. Fifth, manufacturing must be scalable to enable high-throughput manufacturing. Sixth, the setup should require minimal training and utilize readily available raw materials and equipment. Seventh, manufacturing must be simple and require minimal training for expanding production. Finally, there must be easy availability of raw materials and equipment, as well as relative independence from regional utility service fluctuation for production^8^.

To address these challenges, we report the development of **su**rface-enhanced **Ra**man-scattering based **k**it for **s**creening in **h**igh-risk **a**reas (**Suraksha**, Sanskrit for protection), designed to meet the stringent requirements of scalability, cost-effectiveness, and reliability. Our approach relies on affordable, paper-based substrates embedded with dendritic silver nanostructures, which feature a dense array of nanogaps and nanotips to markedly boost the SERS signal. By using easily accessible chemicals, basic electrochemistry techniques, and household equipment, we eliminate the need for cleanroom facilities or advanced laboratory setups. Moreover, a simple battery powers the entire fabrication process, circumventing dependence on an unstable power grid. We validated the reproducibility and robustness of the substrates through rigorous intra-chip, inter-chip, and inter-batch testing, demonstrating consistent plasmonic enhancement.

Furthermore, to demonstrate Suraksha’s real-world effectiveness, we validated it against two critical classes of contaminants, pesticides dissolved in water and bacterial pathogens. In food safety, monitoring pesticide residues is vital to ensure health and regulatory compliance, yet conventional methods such as chromatography and mass spectrometry are expensive, time-intensive, and often impractical for field use. While colorimetric assays and other portable test kits have been developed to address some of these challenges, they can fall short in sensitivity and specificity^9^. SERS offers a compelling alternative, providing rapid, on-site detection of pesticides at relevant concentrations. Its multiplexing ability also makes SERS valuable in environmental monitoring, where agricultural runoff may contain mixtures of pesticides and fertilizers^10–12^. With Suraksha, we achieved a limit of detection (LoD) of 0.1 ppm each for pesticide screening, at an estimated cost of consumables of 1.33¢ per test, making near-instantaneous, on-site diagnostics feasible in resource-limited settings. Similarly, we detect and classify isolates of *Escherichia coli* (E. coli) and *Bacillus subtilis* (B. subtilis) as model bacterial strains. In comparison, conventional methods are not only more expensive, but also require controlled environments, specialized reagents and trained personnel, making them impractical for urgent or resource-limited situations. Collectively, these findings underscore Suraksha’s scalable, accessible, and actionable design, bridging the gap between advanced analytical techniques and point-of-care applications, and empowering communities to reduce cost, complexity, and infrastructure requirements in large-scale screening.

## 2. Results and discussion

### 2.1. Design and development of the *Suraksha* platform

#### 2.1.1. SERS substrate selection and fabrication

In designing a truly field-ready SERS platform, we first ruled out conventional nanopatterning methods that necessitate a clean room facility. These include photolithography, electron beam lithography, nanosphere lithography, focused ion beam technology, chemical vapor deposition, thermal evaporation, and electron beam evaporation, among others^13,14^. Additionally, we also excluded methods requiring specialty chemicals and equipment, controlled temperature or pressure conditions, or inert gas environments. Such methods are not only challenging to implement in resource-limited settings but also prohibitively expensive for commercial scalability^15,16^. Given these constraints, dendritic silver dendritic nanostructures emerged as a viable solution as they can be synthesized in solution and are also known to yield high enhancement factors^17,18^. Among the various fabrication methods for such substrates, sonoelectrochemical^19,20^, photochemical^21,22^, and electroless methods^23–26^ are unsuitable due to the outlined requirements. Similarly, galvanic displacement is disqualified due to the additional steps needed to remove the oxide layer and recover the sacrificed metal^27^. To overcome these limitations, we developed a method for fabricating electrodeposited dendritic silver substrates using inexpensive, commercially available supplies.

Building on conventional electrodeposition methods, briefly described in the Materials & Methods section, we introduced field-friendly modifications^28–30^. A simple battery powered the deposition within a mason jar, used as the electrochemical cell. Instead of expensive gold coated substrates or specialized silver / platinum electrodes, two pieces of Aluminum foil were used as both the working and counter electrodes. Either end of the battery was connected to crocodile clips, via a switch and resistor in one arm. Fig.1 shows the complete battery-powered assembly, including the mason jar setup and 3D-printed jar screw cap replacement, illustrating how the circuit components were spatially integrated into the lid. All connections were made using solder free heat shrink wire connectors to avoid the need for a soldering machine. For large-scale production, the 3D-printed parts can be replaced with injection molded plastic in bulk and included in the kit. All parts used in the fabrication of the kit are readily available on standard online vendors or in a hardware store and do not require access to the power grid to run [Supplementary Table 2].

**Figure 1:**
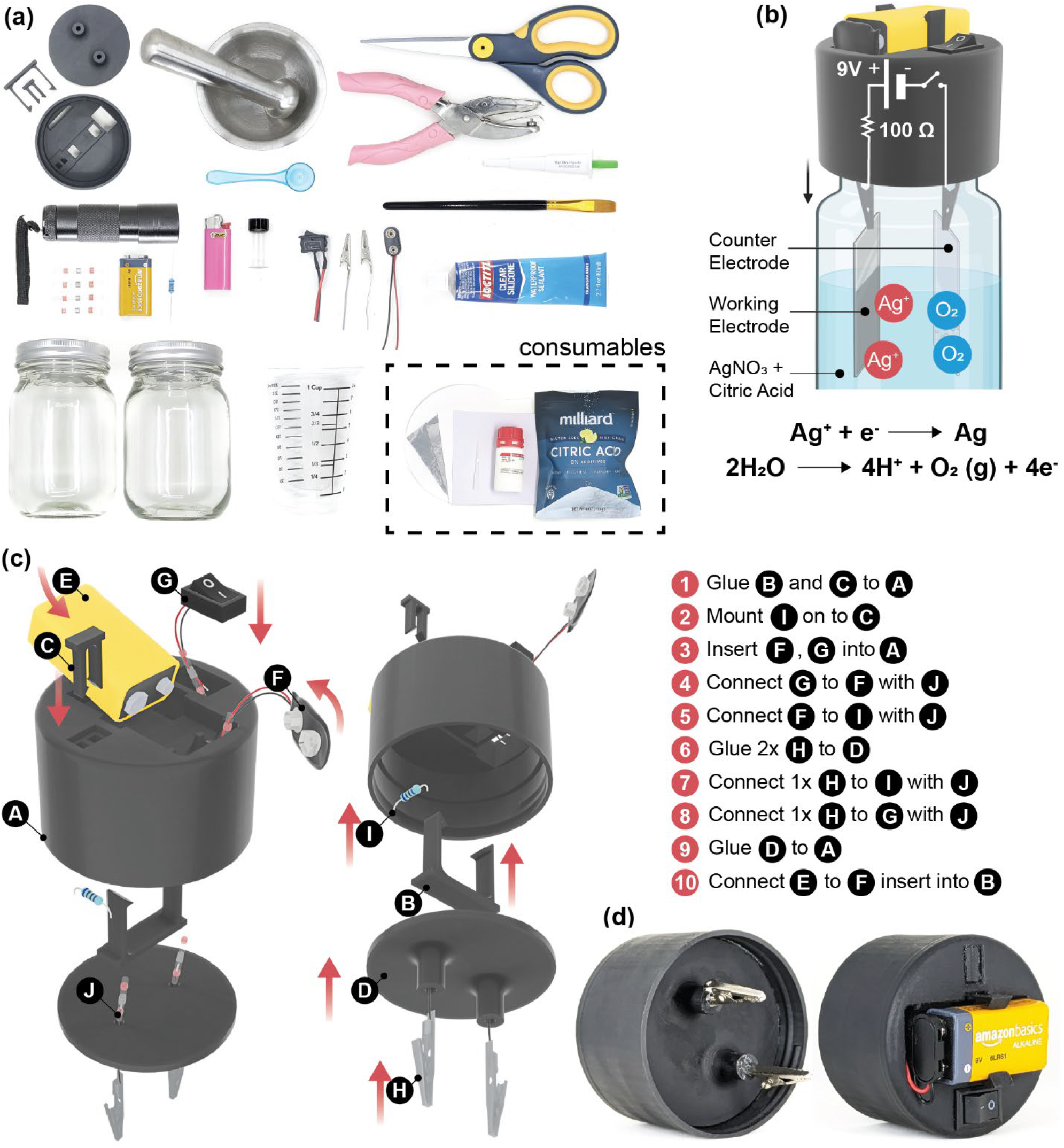
Design of Suraksha kit. (a) Photograph of the components of the Suraksha kit with (inset) consumables used in test fabrication (b) Schematic illustration of the electrochemical cell (c) schematic of the kit assembly of and labelled parts: A: deposition cell, B: battery mount, C: resistor mount, D: electrode mount, E: battery, F: battery connector, G: switch, H: crocodile clips, I: resistor, J: self-solder connector, (d) Photograph of the fully assembled replacement lid of the electrochemical cell

To prepare the electrolyte in resource-limited settings, we simply dissolved 1.5 cups of food-grade citric acid and 0.5 teaspoons of silver nitrate, which are both commonly available reagents, into 5 liters of chlorine-free water, using culinary measurements to avoid the need for a weighing balance. Silver nitrate reacts with dissolved ions like Cl^-^ in tap water to form AgCl, an insoluble white precipitate. However, such precipitates are not observed in distilled water or reverse osmosis (RO) purified water. Many popular drinking water brands sell RO water, and we found that bottled water products that are not remineralized after purification, which typically involves adding some chloride salts, can be used for dendrite fabrication [Supplementary Table 1].

To fabricate the silver dendrites, the aluminum foil electrodes were connected to the crocodile clips, the assembly was screwed into the jar. When the circuit was activated, silver dendrites were deposited on the working electrode. As silver has poor adhesion to the foil due to differences in crystal lattice structures, chemical affinity, and surface smoothness relative to traditional gold-coated substrates^31^, the deposited silver is shed off readily once the battery is turned off. The deposition was repeated cyclically, with intermittent gentle shaking until sufficient nanostructures were formed [Fig. 2a]. In conventional fabrication, where the silver plate’s surface is used as a SERS substrate directly, coverage of the working electrode would be a concern, but as we are collecting the nanoparticles as a powder, even low-cost electrodes and non-uniform depositions prove sufficient. After enough dendrites were generated, the dendrites were collected, filtered, dried, crushed, and resuspended to an optimized concentration for uniform application on our paper-based biosensor.

**Figure 2:**
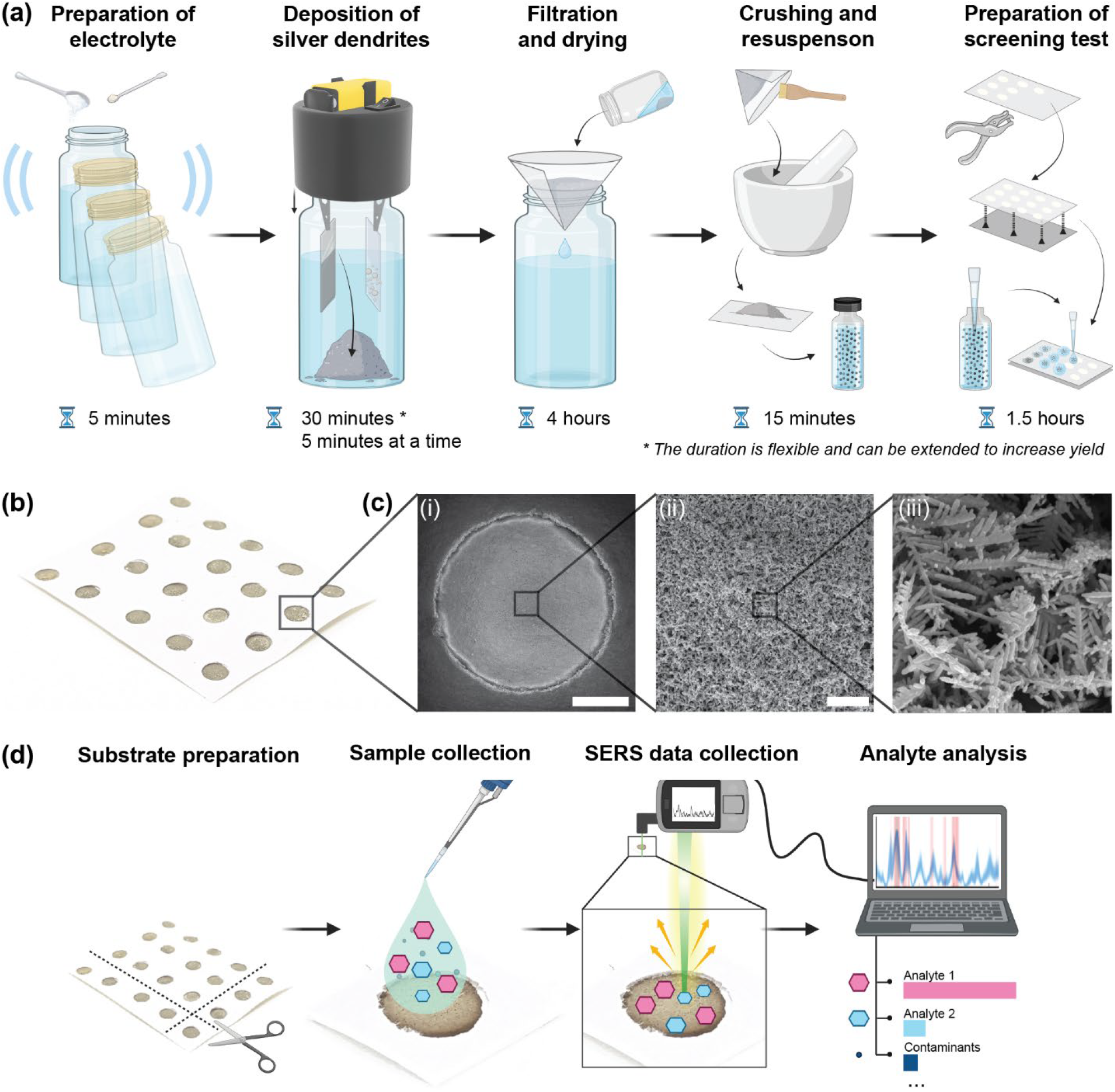
Fabrication of the silver dendrites and assembly of the paper-based SERS substrate. Schematic showing the fabrication of the SERS substrates. (b) Photograph of a strip of fabricated SERS substrates, where each circular region constitutes an individual test. (c) SEM micrographs showing the dendritic nanostructures at three magnifications, with scale bars of i) 1 mm, ii) 5 μm, iii) 1 μm, (d) Schematic representation illustrating how the fabricated tests are used for pesticide detection.

#### 2.1.2. Assembly of the paper-based SERS substrate

To create a robust paper-based SERS chip, we first selected cardstock as the substrate due to its higher GSM (thickness), which prevents sogginess and preserves the integrity of the sensor once silver dendrites are deposited. Because background fluorescence can interfere with SERS measurements^32,33^, choosing low fluorescence paper is crucial. Many papers, particularly those labeled “bright white,” contain optical brightening agents (OBAs) that increase fluorescence background, so a UV flashlight is included in the Suraksha kit to quickly identify and exclude such paper in the field. Under UV light, papers of the same color, with higher fluorescence appear brighter, enabling quick visual comparisons. As demonstrated Fig. 3, while nanomaterial surface energy transfer (NSET) can partially mitigate this fluorescence, it may still impact the overall sensing performance if the correct base paper is not chosen.

**Figure 3:**
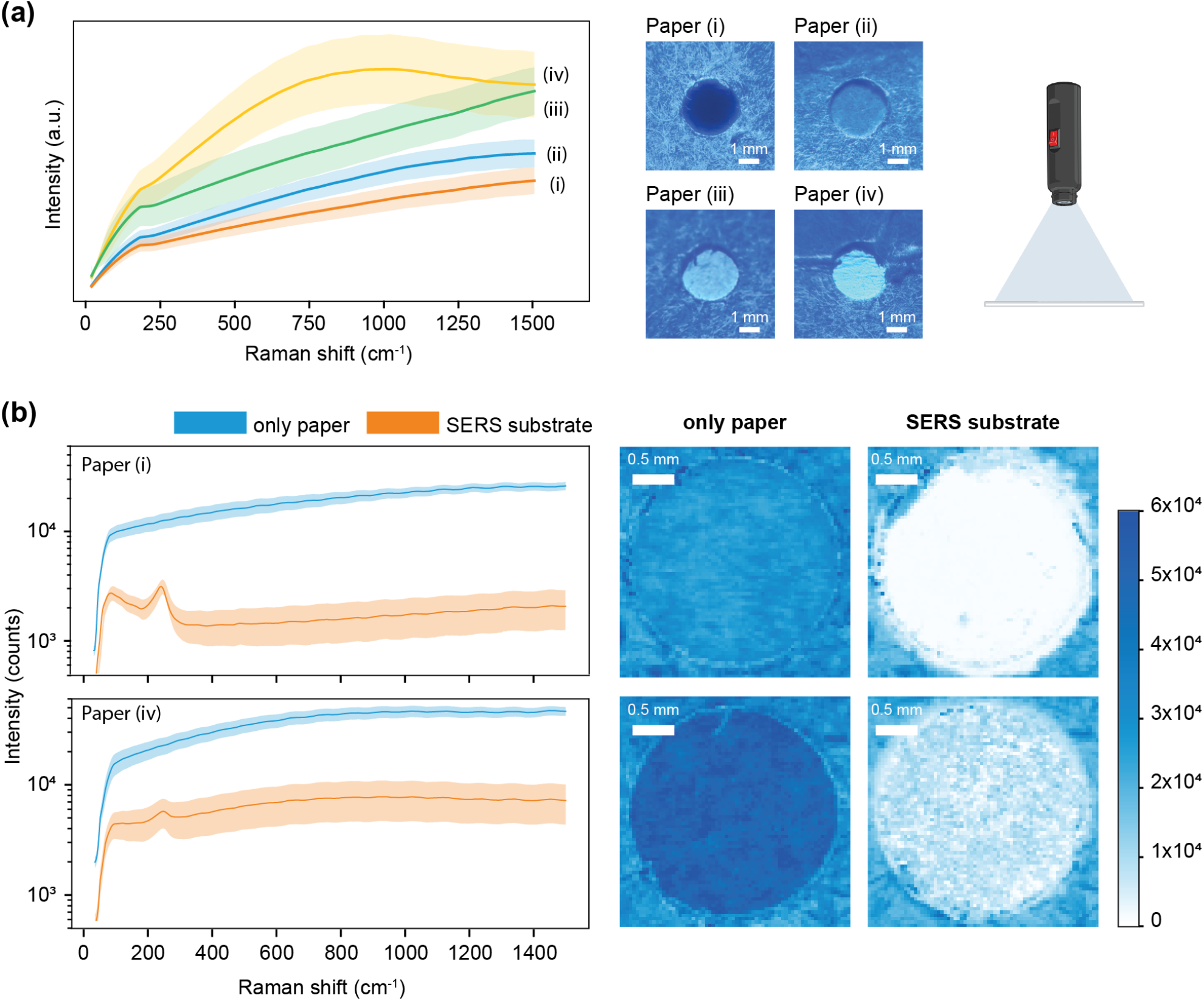
Evaluation of different card stock papers to identify the most suitable option with minimal background fluorescence. (a) Mean Raman spectra (± standard deviation) for four different papers; right panels show UV-illuminated images illustrating background fluorescence. (b) Comparison of the highest and lowest fluorescence paper under two conditions: with and without deposited dendrites, demonstrating the effectiveness of quenching (top: sufficient, bottom: insufficient) for paper via nanomaterial surface energy transfer (NSET). High-resolution Raman maps highlight the quasi-Rayleigh peak intensity, highlight the differences in fluorescence quenching.

For chip fabrication, we punched a grid of holes in hydrophobic paper tape and affixed this grid onto the paper. Each hole was then drop-coated with 10 µL of the colloidal dendrite mixture and allowed to dry. To ensure effective quenching through NSET, it is crucial to achieve both adequate dendrite coverage and uniform distribution. This requires careful dilution and thorough mixing of the colloidal dendrite solution^34^. Fig. 2b shows the completed substrate, while Fig. 2c SEM micrographs reveal densely packed dendritic structures with consistent coverage throughout the punched area. Although the crushing process slightly reduces branching compared to dendrites deposited directly on conductive surfaces, the overall hotspot density remains sufficient for high-quality SERS detection. A comprehensive list of consumables appears in Supplementary Table 3, and detailed protocols are provided in the Materials & Methods section.

### 2.2. Characterization and assessment of designed SERS substrate efficacy

#### 2.2.1. Characterization of optical performance of the substrate

Effective SERS-based detection involves choosing the optimal laser wavelength and power density to maximize the Raman enhancement while minimizing background noise and fluorescence. The degree of enhancement derives from the overlap of the excitation wavelength, the scattering wavelength, and the surface plasmon resonance of the substrate. Additional factors include the analyte’s absorption and the overall optical response of the instrument, encompassing detector properties and optical path absorption^35,36^. To explore this overlap, we investigated the extinction spectrum of the paper-based dendritic substrate, which exhibited a broad peak at ∼315 nm followed by an extended tail across the visible range [Supplementary Fig. 1]^37^. This response is attributed to the nanogap distribution resulting from fractal nanoparticle aggregation on the substrate surface^38^. In selecting the laser power density, we balanced the need for a sufficient SERS signal against avoiding substrate or analyte damage. A power density of 45 mW/cm^2^ at 532 nm was found to be optimal, providing consistent and reproducible signals without impairing the stability of the paper-based sensor [Supplementary Fig. 2]. This combination of moderate laser power and a 532 nm excitation source was adopted for subsequent SERS measurements. Nevertheless, the substrate’s broadband response [Supplementary Fig. 1] permits the use of alternative laser wavelengths for specialized applications, thus extending the versatility of the sensor.

#### 2.2.2. Analyte-independent analysis of substrate using the quasi-Rayleigh peak

To characterize plasmonic enhancement independently of any specific reporter molecule, we utilized quasi-Rayleigh peaks^39–41^ [Supplementary Fig. 3]. These low-frequency scattering peaks occur near the Rayleigh line in Raman spectrometers, arising due to the edge or notch filters that block elastic scattering from the light source. This allows us to unlink the electromagnetic enhancement from the chemical enhancement or interference from the reporter molecule in the SERS signal. It can also serve as an effective quality control strategy for testing substrate enhancement performance without consuming the substrate with a reporter molecule. For our system, quasi-Rayleigh peak measurements at multiple wavelengths revealed that the 532 nm laser provided the most distinguishable signal for evaluating substrate enhancement [Supplementary Fig. 1]. Using this approach, we could confirm the presence of robust localized surface plasmon resonances across the substrate while reserving the option to switch excitation wavelengths for specific target analytes. This quasi-Rayleigh technique thus offers a practical quality control step in resource-limited or field settings by allowing rapid screening of substrate performance before analyte testing.

#### 2.2.3 Optimization of electrodeposition fabrication parameters

Electrodeposition parameters are typically optimized for substrate performance and reproducibility. Cheng et al.’ established a benchmark current density of 1 mA/cm^2^ for silver dendrite morphology^28^, and Amin et al. showed that voltages from 2V to 20V can produce dendrites with varying hotspot densities^42^. However, differences in power supplies, electrode material, and post-deposition treatment necessitate recharacterization. Sensitivity of plasmonic enhancement to deposition voltage is also undesirable because non-standard resistances are hard to find, aluminum foil inconsistencies can alter current density, and battery voltage naturally drops over time (e.g., from 9V to ∼6V before failing)^43^ Substrates fabricated at deposition voltages ranging from 1.5V to 9V were studied via quasi-Rayleigh peak intensities, revealing no significant correlation between plasmonic enhancement and voltage (PCC = 0.17, p = 0.78) [Supplementary Fig. 4]^39–41^. This voltage independence likely stems from post-processing (e.g., crushing the dendrites) and the high density of hotspots on the substrate. The result supports the goal of field deployment, where imprecise manufacturing is expected. For applications requiring a stable battery output, a series of 1.5V silver oxide batteries offers a suitable alternative^44^.

#### 2.2.4. Benchmarking of substrate sensitivity

To assess the sensitivity of the paper-based substrate, we employed methylene blue (MB) as a model Raman reporter. Serial dilutions of MB solutions were prepared, and 20 μL aliquots were drop-cast onto the substrate and allowed to dry. SERS spectra were acquired using a 532 nm laser through a 5× objective lens, with an exposure time of 0.1 seconds without averaging. This configuration enabled high-resolution mapping of 6400 pixels with a spatial step size of 50 μm on each chip, completed within 30 minutes. For rapid screening, fewer points can be measured, reducing acquisition times to under 10 minutes per chip. We processed the spectral maps with a digital SERS protocol that flags a “positive” pixel when the 1624 cm^−1^ MB peak intensity exceeds the local background by at least four standard deviations^45^. This peak-based detection strategy yielded a limit of detection (LoD) between ∼10 nM and ∼100 nM [Fig. 4]. The high-density dendritic features, combined with digital SERS analysis, underscore the sensor’s potential for trace analyte detection in practical scenarios where rapid and sensitive measurements are paramount.

**Figure 4:**
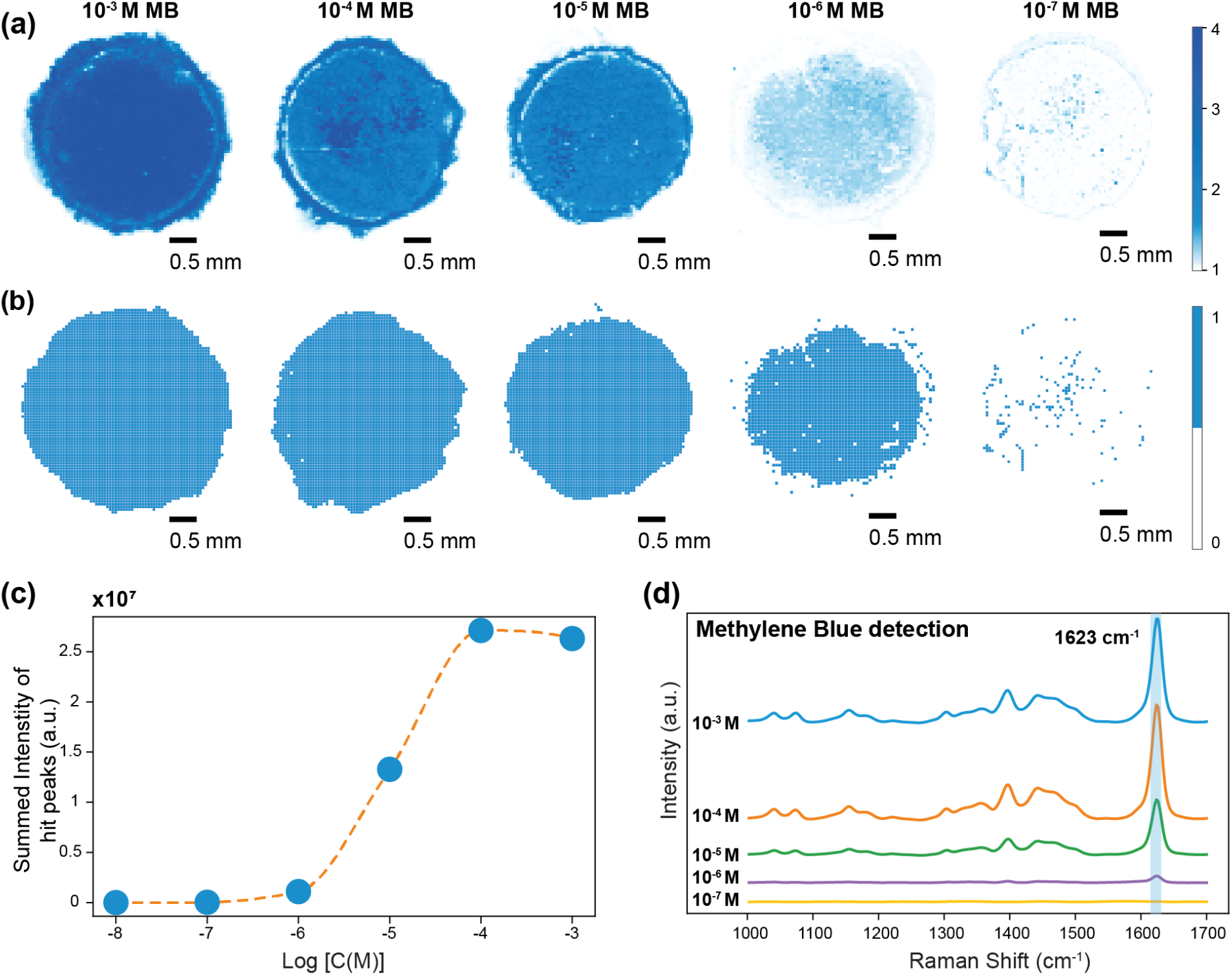
Characterization of the sensitivity of the paper-based silver dendrite biosensor using the standard Raman reporter dye MB across a concentration range of 10^−3^ M to 10^−8^ M. (a): High-resolution intensity map (step size = 50 μm) showing the spatial distribution of Raman signal intensity for MB across an entire chip (1/8” diameter). (b): Digitized intensity maps based on spectral ‘hits’ for MB as determined by the digital SERS method. Map for 10^−8^ omitted for conciseness. (c): Quantitative SERS analysis of MB as a function of concentration, represented by the summed peak intensity at spots with positive signal detections (‘hits’) corresponding to the characteristic Raman peak of MB. (d): Mean spectral plots for MB demonstrating the evolution of the characteristic Raman peaks with concentration.

#### 2.2.5. Validation of plasmonic enhancement reproducibility

Reproducibility is paramount for quantitative and reliable SERS-based sensing, as variability may stem from substrate heterogeneity, batch-to-batch differences, analyte distribution inconsistencies and local electric field variations at plasmonic hotspots^46,47^. To verify the performance, we evaluated the reproducibility of plasmonic enhancement at three levels: (i) intra-chip (different points within a single substrate), (ii) inter-chip (multiple chips within a single fabrication batch), and (iii) inter-batch (chips produced in separate manufacturing runs). To ensure uniform analyte distribution, the substrates were drop-cast with 20 μL of 1 mM MB stock solution and allowed to dry. A high-resolution Raman map was generated with a 532 nm excitation laser and a 5x objective lens across the surface of each chip providing a comprehensive dataset for analysis. Nine chips were prepared (three batches of three chips each), all drop-cast with 20 µL of 1 mM MB to ensure uniform analyte coverage. We then recorded high-resolution Raman maps at 532 nm through a 5× objective lens, processed spectra to remove outliers with Z-scores > 4, and calculated the relative standard deviation (RSD%) for select MB peaks (85 cm^−1^, 450 cm^−1^, 770 cm^−1^, and 1624 cm^−1^). The 85 cm^−1^ quasi-Rayleigh peak, reflecting plasmonic enhancement independent of analyte distribution, served as a direct measure of substrate uniformity, while the other peaks, corresponding to vibrational modes of MB, reinforced the findings^39–41^ [Fig. 5, Supplementary Fig. 5]. Low RSD% values were observed across all levels of variability, with even inter-batch RSD% across all batches consistently below 10%. The contour plot in Fig. 5e further emphasizes this reproducibility across the dataset by demonstrating the consistency in intensity across the Raman Shift range.

**Figure 5:**
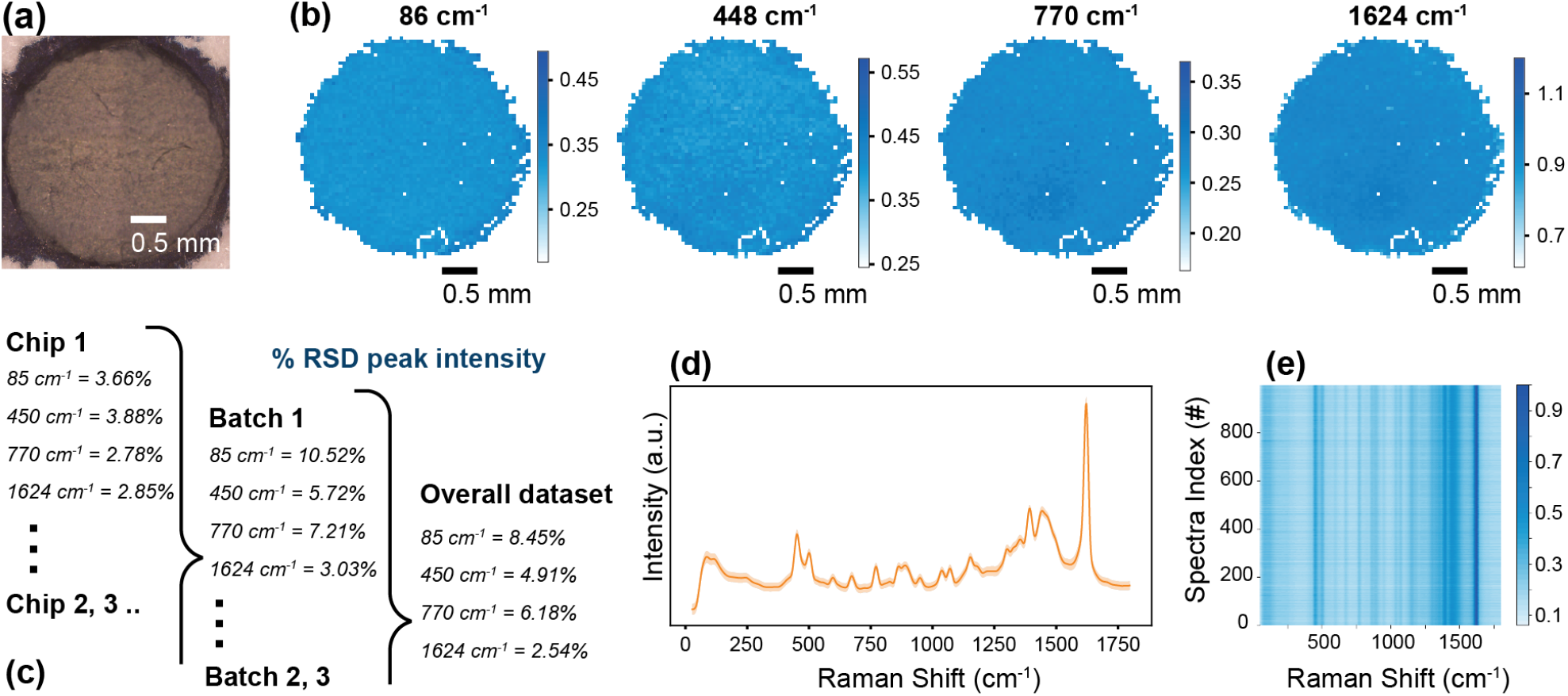
Reproducibility of the paper-based silver dendrite substrates. (a) Optical microscope image of a representative sensor chip. (b) High-resolution Raman mapping of MB at 86, 448, 770, and 1624 cm^−1^; the 86 cm^−1^ quasi-Rayleigh peak signifies substrate enhancement. (c) RSD of the peak intensities showing intra-chip (one chip), inter-chip (three chips in one batch), and inter-batch (three batches of three chips each) variability. (d) Mean Raman spectrum (± standard deviation) across the full dataset, reflecting consistency in spectral features. (e) Contour plot of 1000 randomly sampled spectra from the dataset, indicating reproducibility.

Notably, the paper in the substrates effectively wicks the liquid sample during drying, preventing coffee ring formation, a common problem on rigid glass or metal supports^48,49^. By preventing analyte aggregation at the droplet perimeter, these substrates yield more homogeneous SERS signals and improved point-to-point reproducibility. Supplementary Fig. 6 illustrates the severity of coffee-ring patterns on typical rigid substrates compared to the uniform distribution on our paper-based sensors. Additionally, on these conventional substrates, the spreading shape of the analyte is not controlled and can become irregular due to surface interactions. Consequently, this paper-based platform promises robust, large-scale SERS analysis with minimal positional variability—an essential attribute for reliable field testing and high-throughput screening.

## 3. Demonstration of field applications of the biosensor

### 3.1. Pesticide detection

Growing concerns over pesticide residues in agricultural runoff necessitate highly sensitive and field-viable analytical methods for ensuring environmental and food safety. The Environmental Protection Agency (EPA) has established guidelines defining permissible levels of pesticides in various food products^50^. While traditional laboratory-grade tools like gas chromatography, high-performance liquid chromatography (HPLC), chromatography-mass spectrometry (GCMS) offer high sensitivity, they often face challenges, including high costs, bulkiness, lack of portability, need for trained personnel and time taken for detection. Existing field-based tests, such as colorimetric assays, are more convenient but lack the sensitivity needed for low-concentration detection, typically operating in the range of 1–10 ppm. Cross-reactivity can lead to false positives or negatives, especially when testing complex samples containing multiple pesticides or related compounds and reagents need to remain stable under varying conditions, such as high temperatures or humidity^9^.

In this study, we investigated the label-free SERS-based detection of two pesticides, thiram and thiabendazole (Fig. 6). Using our silver dendrite substrates, we observed the highest signal enhancement at a 785 nm excitation wavelength. Serial dilutions of thiram and thiabendazole were prepared to assess both sensitivity and quantitative performance. A 20 μL aliquot of each dilution was drop-cast onto the substrate and dried at ambient conditions. The acquired spectra were pre-processed and analyzed via digital SERS algorithms focusing on characteristic Raman peaks at 1383 cm^−1^ (thiram) and 785 cm^−1^ (thiabendazole). To further explore specificity and multiplexing capabilities, we conducted additional experiments using a mixture of both pesticides. We generated combined digital SERS maps, labeling pixels based on whether they contained thiram, thiabendazole, neither, or both. This multiplexing feature is highly relevant in agricultural runoff, where multiple pesticide residues and fertilizers often coexist. The results showed that both pesticides were individually detectable down to 0.1 ppm, and the multiplexed detection maintained the same sensitivity (Fig. 6b(iii). These findings are comparable to other label-free SERS methods employing silver dendrites and lab-grade substrates. Notably, the ease of use and field compatibility of our substrates underscore their potential for on-site pesticide monitoring. Moreover, the successful demonstration of simultaneous detection of multiple analytes highlights the versatility of this approach for broader applications in environmental diagnostics.

**Figure 6:**
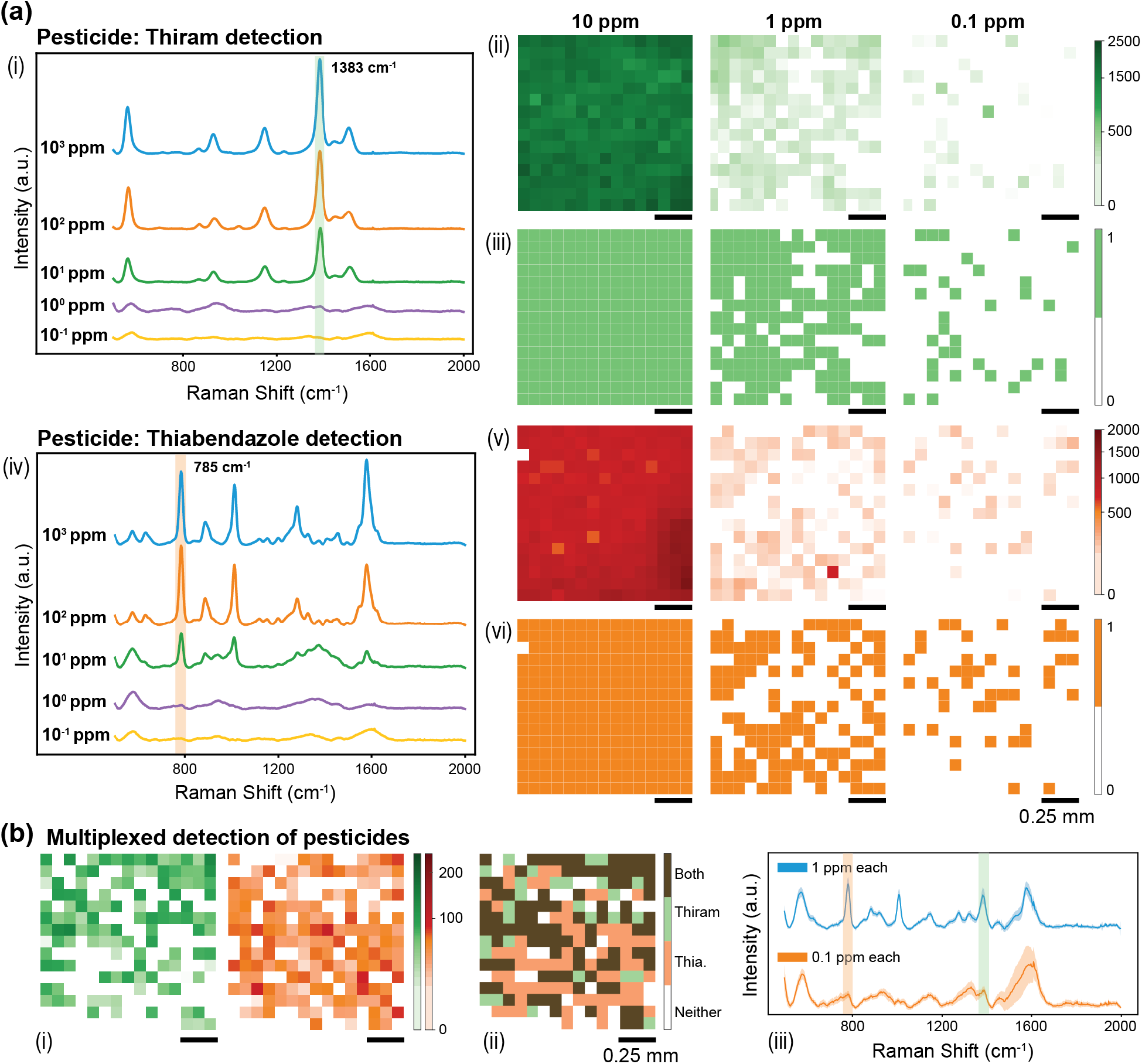
Pesticide detection using the paper-based SERS substrate. (a) Detection of thiram (i– iii) and thiabendazole (iv–vi) from 10^3^ to 10^−1^ ppm (the 10^2^, 10^1^ ppm maps are omitted for space), targeting characteristic Raman peaks at 1383 cm^−1^ (thiram) and 785 cm^−1^ (thiabendazole). (i, iv) Mean spectra illustrating peak evolution with concentration. (ii, v) Intensity maps of the 15×15 grid (1.25×1.25 mm) showing spatial signal distribution. (iii, vi) Digitized maps using the digital SERS method to identify spectral “hits.” Additional quantitative results appear in Supplementary Fig. 7.(b) Multiplexed detection of thiram and thiabendazole at 100 and 10^−1^ ppm: (i) Individual intensity maps highlighting detection regions for each pesticide (the 10^1^ ppm map is omitted). (ii) Combined heatmap showing co-detection (brown), thiram-only (green), thiabendazole-only (orange), and no detection (white). (iii) Mean spectra for both pesticides, displaying their respective peaks at 1383 cm^−1^ (thiram) and 785 cm^−1^ (thiabendazole). Supplementary Fig. 8 provides additional digitized maps.

**Figure 7:**
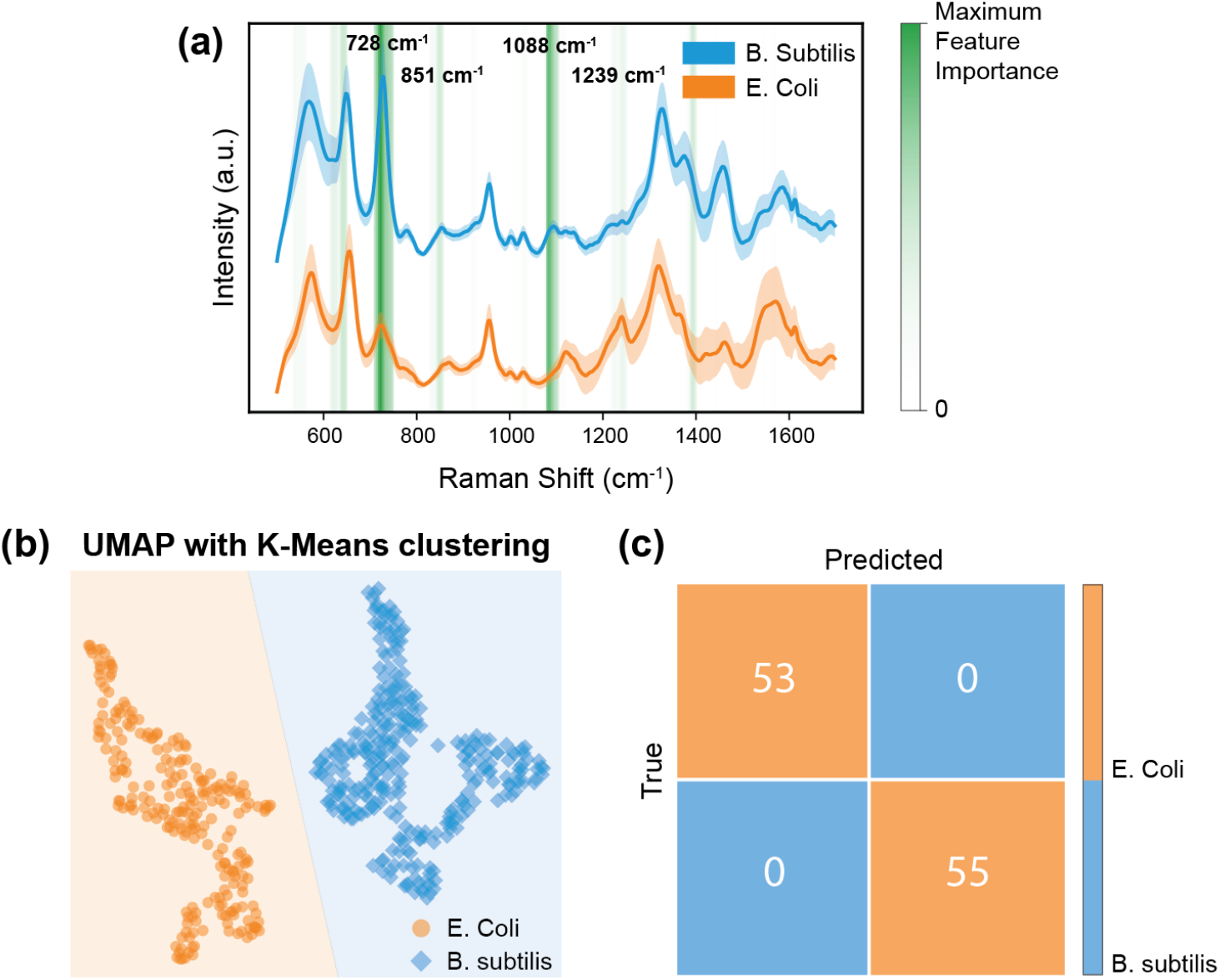
Bacteria classification using the developed SERS substrate. (a) Mean Raman spectra of B. subtilis (blue) and E. coli (orange), highlighting key discriminatory peaks (green), derived from random forest feature importance analysis. (b) UMAP projection of the spectra, followed by k-Means clustering, illustrating distinct separation between the two bacterial species. (c) Confusion matrix for the random classifier on a 20% held-out test set, confirming high accuracy in distinguishing B. subtilis from E. coli using SERS signatures

### 3.2. Bacteria classification

Rapid and accurate detection of bacterial pathogens is critical for managing public health risks, especially in resource-limited settings. Traditional methods such as polymerase chain reaction (PCR) and Matrix-Assisted Laser Desorption/Ionization Time-of-Flight (MALDI TOF) are expensive, require specialized reagents, trained personnel, controlled environments and laboratory facilities, making them unsuitable for point-of-care deployment. Label-free SERS combined with machine learning and a handheld spectrometer provides a rapid, portable, and reagent-free platform that further enhances field applicability^51^.

We used the SERS substrate for demonstrating bacterial discrimination using E. coli and B. subtilis as model pathogens. Bacterial cultures were washed to eliminate spectral contaminants, centrifuged to form a pellet, and resuspended in deionized water. Following a drop-cast of 20 µL onto the silver dendrite substrates and drying under ambient conditions, SERS spectra were collected on a 10×10 grid for each species. To reduce dimensionality while preserving spectral relationships, we employed Uniform Manifold Approximation and Projection (UMAP), followed by k-Means clustering to group the spectra into two distinct clusters. Separately, a Random Forest classifier was also trained on the dataset, achieving 100% accuracy on a 20% held-out test set, indicating excellent performance under ideal experimental conditions. Each decision tree in the ensemble made splits based on the Gini impurity, which quantifies how effectively a split separates different classes. A feature importance analysis, based on the Gini importance metric, revealed that the Raman shifts at 728 cm^−1^ (adenine ring breathing mode, nucleic acids), 851 cm^−1^ (C–C stretching in proteins), 1088 cm^−1^ (phosphate backbone of nucleic acids), and 1239 cm^−1^ (amide III band from proteins/peptides) played crucial roles in discriminating bacterial species^52^.

These results demonstrate that, once bacteria are appropriately isolated in the field and resuspended at optimal concentrations, they can be reliably identified on-site using machine learning-assisted, label-free SERS. This study serves as a proof of concept for a scalable, high-specificity platform for rapid bacterial screening.

## 4. Cost and logistics of substrate fabrication in the field

The primary goal of this study was to develop an affordable and easily scalable SERS platform to meet the demands of decentralized testing. The entire kit assembly requires 15–30 minutes of operator time, followed by a drying period, ensuring minimal training barriers for resource-constrained environments. At retail prices in the USA from standard online vendors, the total cost of assembling one kit is $39.54 [see Supplementary Table 2], making this approach highly economical and feasible for large-scale deployment. Each mason jar deposition yields approximately 42.5 mg of silver dendrites in 30 minutes, sufficient for ∼425 tests at a preparation concentration of 10 mg/mL. The cost per test thus approximates 1.33¢ [see Supplementary Table 1], a remarkably low figure that highlights the cost-efficiency of this method. Such affordability positions this process as a practical solution for widespread testing, particularly in low- and middle-income countries (LMICs) where cost constraints often limit access to advanced screening and monitoring tools.

The overall fabrication process for a batch of substrates can be completed within seven hours, with only ∼90 minutes of hands-on labor [Fig. 2d]. This efficiency allows operators to work on multiple kits in parallel, optimizing throughput. For instance, to test 10$ of the population of a town with 50,000 people daily, mirroring testing demands observed during peak pandemic conditions, only four staff members operating in parallel would be required assuming a standard 8-hour workday with a 1-hour lunch break^53^. One technician operates five testing kits to fabricate the silver nanostructures, another filters the silver dendrites, one grinds the silver to a fine powder with a mortar and pestle to form the colloid and makes the paper grid, while the last drop the colloid onto grids to make the final substrate. Such a modular framework enables each site to easily scale production by adding more operators or additional kits.

## 5. Conclusions

In conclusion, the Suraksha platform presents an accessible, cost-effective and highly adaptable solution for decentralized biosensing, with applications encompassing environmental monitoring, food safety, and public health. By introducing a SERS substrate fabrication kit that enables rapid, large-scale production, we lower the barriers to adoption of this technology and empower small to medium-sized towns and resource-limited areas to become self-reliant in addressing critical testing needs. We demonstrated high sensitivity and multiplexing in pesticide detection, revealing limits of detection of approximately 0.1 ppm for thiram and thiabendazole, levels that surpass many conventional field-based colorimetric tests. We also showcased robust bacterial classification for *E. Coli* and *B. Subtilis* when paired with machine-learning algorithms, evidencing the broad utility of our label-free SERS platform for pathogen screening and beyond. Validation studies using MB confirmed the reproducibility of the test substrate, with minimal variation across intra-chip, inter-chip, and inter-batch measurements.

Our analysis addresses critical economic, supply chain, and design considerations to ensure ease of fabrication and scalability. The cost of producing the fabrication kit is $39.54 with raw materials purchased at bulk retail prices, while the consumable cost for fabricating each test is as low as 1.33¢. Minimal sample preparation and no specialized expertise make it ideal for on-site testing with a portable Raman spectrometer. The label-free nature of our detection method circumvents bottlenecks associated with reagent development, facilitating the rapid adaptation of the platform to new targets, such as emerging pollutants or novel pathogen. Furthermore, the digital integration capabilities of SERS enable a ‘hub and spoke’ digital integration, allowing newly detected compound spectra to be shared across field spectrometers. Overall, our platform’s affordability, adaptability, and scalability set a new standard for decentralized testing, representing a significant stride toward inclusive and reliable biosensing across diverse and underserved settings worldwide.

## 6. Experimental section

### 6.1 Materials

Methylene blue (MB), Thiram, and Thiabendazole were purchased from Sigma Aldrich (USA). Escherichia coli DH5α and Bacillus subtilis strains were obtained from the American Type Culture Collection (ATCC, USA). Luria–Bertani (LB) agar plates were purchased from ThermoFisher Scientific (USA). Detailed lists of additional chemicals, reagents, and equipment used for the fabrication of the Suraksha kit, as well as the consumable tests and recommended drinking water brands, are provided in Supplementary Tables 1–3.

### 6.2. Methods

#### 6.2.1. Fabrication of standard silver dendrite SERS substrate for reference

A single-step electrochemical deposition technique was employed to fabricate nanostructures on gold-coated glass slides^30,54^. The gold-coated substrate served as the working electrode (WE), while a platinum sheet or wire served as the counter electrode (CE). Both electrodes were placed 3 cm apart in an electrolyte containing 2 g/L silver nitrate and 40 g/L citric acid in deionized water. Citric acid selectively binds to certain crystallographic planes of silver, guiding dendritic growth. An AMETEK SI VersaSTAT 3 benchtop potentiostat supplied a constant current density of 1 mA/cm^2^ for 300 s, forming dense silver dendrites via oriented hierarchical attachment. After deposition, the substrates were rinsed with deionized water and dried under a gentle nitrogen stream.

#### 6.2.2. Design and prototyping of the Suraksha kit

A 9V alkaline battery was used to power the electrodeposition in a mason jar, which served as the electrochemical cell. Two pieces of aluminum foil were used as the working (WE) and counter electrodes (CE). A 100-ohm resistor was inserted in series to drop the voltage to approximately 2.8 V, yielding a current density of about 1.12 mA/cm^2^. A simple switch was incorporated to activate or deactivate the circuit. All connections were established using solder-free heat-shrink connectors, sealed with a BIC mini lighter. Either end of the battery was connected to a crocodile clip via the switch and resistor in one arm. A custom 3D-printed holder designed in SolidWorks was prototyped using a Uniformation GKtwo resin printer with Uniformation GKtwo PLA plant-based black resin. The chassis supported the battery, resistor, switch, and crocodile clips with an integrated screw to mount on to the mason jar. The circuitry was separated from the electrolyte using a 3-D printed base which was glued to the rest of the assembly and sealed using Loctite Clear Sealant. The printed ensured isolation from the electrolyte, exposing only the crocodile clip ends to the solution. The parts were designed to keep ease of printing and weight of material in mind.

#### 6.2.3. Fabrication of paper-based plasmonic biosensor

An electrolyte solution containing 40 g/L citric acid and 2 g/L silver nitrate was prepared by adding 1.5 cups of food-grade citric acid and 0.5 teaspoons of silver nitrate to 5 liters of chlorine-free water. The solution was mixed in a mason jar by shaking until no visible clumps remained. The aluminum foil electrodes were then immersed in this solution, and the battery circuit was switched on for 5-minute intervals. During electrodeposition, silver dendrites gradually formed on the working electrode. After each 5-minute deposition step, the nanostructures were dislodged from the working electrode by gentle shaking. The resulting dispersion of dendrites was filtered through Whatman paper and left to dry for at least 4 hours (drying could be accelerated with mild heating). Once dried, the dendrites were crushed with a mortar and pestle to produce a fine powder. A paintbrush facilitated transfer of the powder, minimizing loss. The powder was diluted to roughly 10 mg/mL by dispersing in 1 teaspoon of water (4 mL). A 1/8-inch hole puncher was used to prepare a grid pattern in a strip of hydrophobic paper tape, which was then adhered to cardstock. A mini fixed-volume micropipette delivered 10 µL of the silver dendrite mixture to each hole, forming discrete sample spots. The nanostructure-coated spots were allowed to air dry for about 90 minutes (or faster with mild heating) before further use. To prevent cross-contamination between test areas, the cardstock was pre-cut into individual squares measuring 1/4 inch per side before use for sensing applications.

#### 6.2.4. Optical reflectance characterization of silver dendrite surfaces

A PerkinElmer Lambda 950 UV/VIS/NIR spectrometer was used for reflectance characterization of the substrate. The silver dendrite nanostructures were deposited over a card stock paper comparable in size the output port of the integrating sphere of the spectrometer to ensure complete coverage [see Supplementary Fig. 1].

#### 6.2.5. Assessment of burning power threshold on silver dendrite surfaces

The laser power at the sample was measured using a Genentec EO PRONTO Si power meter and determined to be 66.4 mW at the objective, with a spot size of approximately 43 µm (corresponding to 4500 W/cm^2^ at 100% power at 532 nm laser). Time-resolved Raman spectra under these conditions were recorded with a Horiba XploRa PLUS confocal Raman microscope equipped with a 5× objective to determine acceptable duration of exposure [see Supplementary Fig. 2].

#### 6.2.6. Verification of equivalence between supplied voltage and desired current density

The resistance of the electrochemical cell was experimentally determined using a AMETEK SI VersaSTAT 3 potentiostat. The potentiostat was used in a two-electrode configuration with two aluminum foil pieces (7 cm x 4 cm, both sides conducting) as electrodes. A current density of 1 mA/cm^2^ was supplied in chronopotentiometry mode over 300 seconds, and the potential was recorded to determine the voltage required and calculate the cell resistance as approximately 42.85 ohms. Similarly, a voltage of 2.5V was applied in chronoamperometry mode to verify the achievement of the desired current density.

#### 6.2.7. Characterization of substrate plasmonic enhancement response to deposition voltage

A VersaSTAT3 Potentiostat Galvanostat was used to fabricate substrates at deposition voltages of 1.5V, 2V, 4V, and 9V. A Horiba XploRa PLUS confocal Raman microscope with a 5x objective lens was used to study the relative enhancement of the quasi-Rayleigh peak at an excitation wavelength of 532 nm. Spectra from 3 different substrates at each voltage were recorded with an exposure of 1 second, which were averaged over 5 scans. A Pearson Correlation Coefficient (PCC) was calculated to study the correlation between deposition voltage and the intensity of the quasi Rayleigh peak.

#### 6.2.8. Sample preparation for SERS analysis

A 10 mM stock solution of MB was prepared in deionized water. Subsequent working solutions were obtained by serially diluting the 10 mM stock in water to achieve the desired final concentrations. Stock solutions of thiram and thiabendazole were each prepared at 1000 ppm (mg/L) in methanol, due to low water solubility. For experimental use, these stock solutions were diluted with deionized water to obtain the required working concentrations. Care was taken to ensure thorough mixing at each dilution step, and all prepared solutions were used promptly.

Preceptrol cultures of bacteria used, i.e., E. coli DH5α and B. subtilis were rehydrated and plated on LB agar and incubated at 37 °C and allowed to grow for 24 hours. The bacteria were then harvested and transferred to 1 mL of DI water, centrifuged at 3500 rpm for 4 minutes to purify, and the pellet was resuspended at high concentration in DI water. 20 µL sample volume was immediately spotted on each substrate to prevent lysis of bacteria in hypotonic environment. All culture steps were conducted under aseptic conditions.

#### 6.2.9. SERS measurements on fabricated substrate

All SERS measurements for substrate characterization were performed using the SWIFT mode on a Horiba XploRa PLUS confocal Raman microscope equipped with a 5× objective 1200 cm^-1^ grating, and a 532 nm laser source. Each spectrum was acquired with an exposure time of 0.1 seconds, without signal averaging, enabling rapid data collection. The high-speed mapping mode captured the spectra from 6400 pixels (80×80 grid), with a spatial step size of 50 μm. For pesticide and bacterial samples, 785 nm excitation wavelength showed better results, and was used with a 600 cm^-1^ grating. Spectra were acquired using a 1-second exposure time and each spectrum was averaged several times to minimize noise and enhance the signal-to-noise ratio with a 5x objective lens. All acquired spectral data underwent preprocessing to ensure accuracy and reliability. A Savitzky-Golay filter was applied for high-frequency noise reduction, while baseline correction was performed to remove background fluorescence contributions. Spectra were then trimmed to the desired Raman shift range and normalized, if necessary, to allow for consistent peak-intensity comparisons between samples. To eliminate anomalies, outlier removal was performed by discarding spectra exceeding the desired Z-score threshold.

#### 6.2.10. Digital SERS analysis for pesticide detection

For spectral classification, a digital SERS analysis approach was conducted to determine whether specific Raman peaks exceeded a threshold relative to background noise. The characteristic intensity of a target Raman peak was compared against a background intensity range. A spectrum was classified as positive (or ‘hit’) if its peak intensity exceeded the local average baseline by four standard deviations. This approach allowed for the digitization of Raman maps, enabling spatial representation of the detected analytes across the SERS substrate.

#### 6.2.11. Machine learning analysis for bacteria classification

After data pre-processing, Uniform Manifold Approximation and Projection (UMAP) was used for dimensionality reduction, transforming high-dimensional spectral data into a 2D feature space while preserving local relationships. Following dimensionality reduction, k-Means clustering was applied to the UMAP-reduced dataset to group spectra into 2 clusters corresponding to bacterial species.

For supervised classification, a Random Forest classifier with 50 decision trees was trained on 80% of the dataset, with the remaining 20% used for testing and reduction in Gini impurity was used to separate the bacterial species. Gini impurity is calculated as:

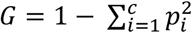, where *pi* represents the proportion of samples belonging to class *i* in a node, and *c* is the total number of classes.

## ACKNOWLEDGEMENT

This work was supported by the National Institute of General Medical Sciences (grant 1R35GM149272). Figures 1b, 2a, 2d, 3a were created using BioRender.com. The authors would also like to thank Prof. David Gracias and Dr. Arijit Ghosh for their input in this project.

