## Supplementary information for "Affordable plasmonic biosensing: democratizing SERS with scalable, field-compatible substrate fabrication"

<u>Supporting information</u>

**Supplementary table 1:** Water testing to determine suitable water for electrodeposition of silver dendrites. In the presence of chloride ions, silver nitrate reacts to form silver chloride, which is a white, insoluble precipitate. A concentrated solution of silver nitrate (20 g/L) was added to 5 mL of each water sample to test for insoluble silver chloride formation. The results suggest reverse osmosis and distillation are effective purification techniques

| **Was precipitation of silver chloride observed on addition of silver nitrate?** | |
|---|---|
| Distilled water | No |
| Reverse osmosis purified water (*ex. Aquafina, 7/11 pure water*) | No |
| Remineralized reverse osmosis purified water (*ex. Essentials*) | Intermediate |
| Boiled water | Intermediate |
| Filtered water | Yes |
| Water treated with Campden tablets | Yes |
| Tap water | Yes |

**Supplementary table 2:** Cost of consumables for each SERS test based on retail prices. The cost is based on the quantity of consumables needed for 425 tests, which are produced from a 30-minute deposition batch using a fresh 9V battery. All prices noted are based on online prices in the USA and are subject to change.

| **Cost of Consumables for Each Fabricated Test (Retail prices)** | **Cost / Test** (0.01 USD) |
|---|---|
| 1. Food grade citric acid (Amazon ASIN: B002DCPR9C) | 0.0323 |
| 2. Silver nitrate (Sigma Aldrich SKU: 204390-2KG) | 1.1341 |
| 3. Distilled water (Amazon ASIN: B0CKLT2TFL) | 0.0870 |
| 4. Whatman filter paper (Amazon ASIN: B00AZ3ARNC) | 0.0188 |
| 5. Aluminum foil (Amazon ASIN: B07B414NZ4) | 0.0000 |
| 6. Cream Card stock paper (Amazon ASIN: B0BWMR7GWM) | 0.0140 |
| 7. Hydrophobic paper tape (Amazon ASIN: B096SF6MDT) | 0.0334 |
| 8. 20 µL pipette tip (Sigma Aldrich SKU: AXYTF20) | 0.0182 |
| **Total cost of each test:** | **1.34¢** |

**Supplementary table 3:** Cost of materials for assembling the Suraksha kit. Listed items are grouped by component category, with retail prices from U.S. online sources dated 15th October 2024. The cost of 3D-printed parts is based on Uniformation GKtwo PLA resin. Whole-sale purchasing or large-scale manufacturing (e.g., injection molding) could significantly reduce costs. Electronics purchased had integrated wires to allow solder free heat shrinks to be used for connection. The resistor was used to stabilize and control the yield of the kit. Metal mortar-and-pestles have lesser losses of powder while crushing compared to porcelain or glass.

| Bill of Materials: Substrate Fabrication Kit (Retail prices) | **Cost / Kit** (USD) | (%) |
|---|---|---|
| ***1. 3-D printed assembly*** | **2.91** | **7.37** |
| • Deposition cell chassis (volume = 84.73 mL) | 2.12 | |
| • Battery mount (volume = 5.09 mL) | 0.13 | |
| • Resistor mount (volume = 24.63 mL) | 0.06 | |
| • Electrode mount (volume = 2.28 mL) | 0.61 | |
| ***2. Electronics*** | **3.61** | **9.13** |
| • 9V alkaline battery (Amazon ASIN: B00BGIVI1K) | 1.42 | |
| • 9V battery connector (Amazon ASIN: B08SL9X2YC) | 0.50 | |
| • Rocker switch (Amazon: ASIN: B07FTXKKZ4) | 0.58 | |
| • 2x Crocodile clips (Amazon ASIN: B07WBWT8CF) | 0.76 | |
| • 4x Heat shrink solder (Amazon ASIN: B087PQNKQR) | 0.31 | |
| • Resistor (100 ohms, Amazon ASIN: B09WJY76F6) | 0.05 | |
| ***3. Labware*** | **4.10** | **10.37** |
| • 2x 16 oz Mason jar (Amazon ASIN: B0CSYW1XB4) | 3.07 | |
| • 5 mL Glass vial (Amazon ASIN: B09GNFVZDB) | 0.26 | |
| • Teaspoon (Amazon ASIN: B09SJT4B9Y) | 0.50 | |
| • Cup measure (Amazon ASIN: B09SGTJCSF) | 0.28 | |
| ***4. Lab Equipment*** | **27.18** | **68.74** |
| • Loctite Clear Sealant (Amazon ASIN: B0002BBX3U) | 5.34 | |
| • BIC Mini Lighter (Amazon ASIN: B077DL93G1) | 0.37 | |
| • Mortar and pestle (Amazon ASIN: B0D7ZW8ZV1) | 13.99 | |
| • Fixed volume pipette (Amazon ASIN: B0BVZ1MHFP) | 7.49 | |

| | | |
|---|---|---|
| ***5. Stationery*** | **1.74** | **4.39** |
| • Scissor (Amazon ASIN: B07RQ9W124) | 0.35 | |
| • Hole punch (1/8 inch, Amazon ASIN: B0B4MKKKLD) | 1.30 | |
| • Paint brush (Flat, Amazon ASIN: B07QN3TV92) | 0.09 | |
| **Total cost of each kit:** | **$39.54** | **100 %** |

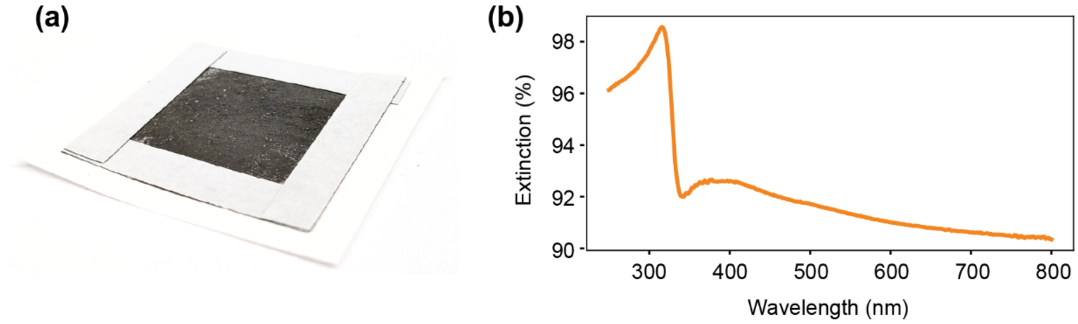


**Supplementary figure 1**: Optical characterization of silver dendritic powder and substrate. (a) Digital image and (b) UV-Vis extinction spectrum of the fabricated substrate.

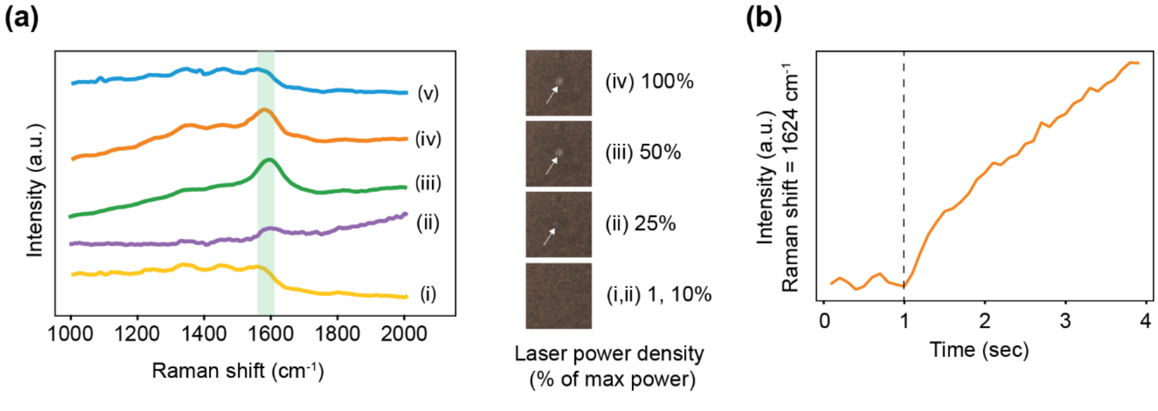


**Supplementary figure 2**: Power density optimization. (a) Representative Raman spectra of the substrate background at 532 nm under varying power densities (1%–100%). At 10% power, an amorphous carbon peak appears near 1600 $cm^{-1}$ without visible substrate damage; at 25% power, both the carbon peak and substrate burning become evident. At ≥50% power, strong fluorescence from the paper substrate obscures the carbon peak. Consequently, 1% power

(45 mW/cm²) was selected to preserve substrate integrity and ensure reproducibility. (b) At 10% power (450 mW/cm²), the carbon peak remains minimal for up to 1 s of exposure, enabling short, high-power scans for rapid data collection before substrate degradation becomes significant.

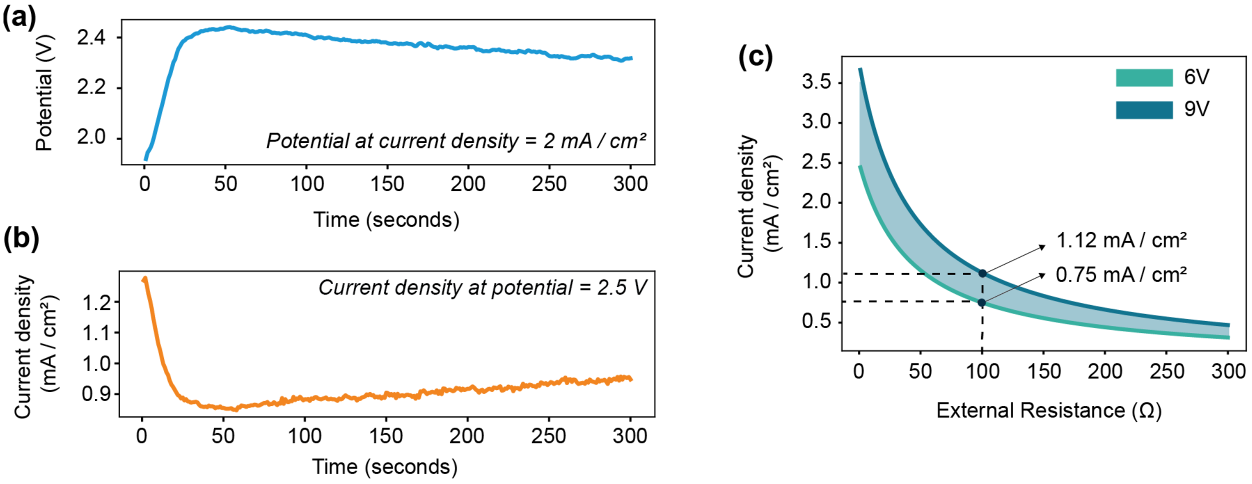


**Supplementary figure 3:** Voltage-Current characteristics of the electrodeposition cell. (a) A current density of 1 mA/cm² was supplied over 300 seconds, and the potential was recorded to determine the voltage required and calculate the cell resistance as $R_{cell} = V_{measured} / i_{applied}$ = 42.85 ohm (approximate). (b) Applying 2.5 V confirmed the feasibility of achieving the desired current density. (c) Estimated operating current density as the battery voltage declines from 9 V to 6 V, where $j = V_{battery} / (A_{electrode} * [R_{cell} + R_{additional}])$.

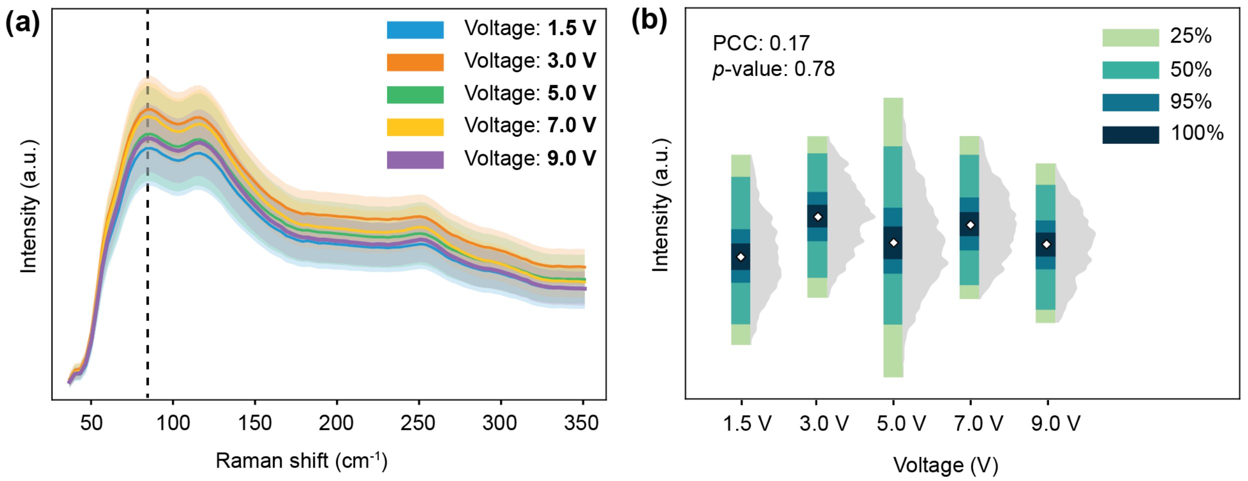


**Supplementary figure 4:** Sensitivity of substrate enhancement performance to deposition voltage. (a) Quasi-Rayleigh peak intensity at an excitation wavelength of 532 nm from three different substrates at each deposition voltage, averaged over five 1s scans. (b) Peak intensity plotted against deposition voltage yielded a Pearson correlation coefficient of -0.17 ($p = 0.78$), indicating no statistically significant relationship.

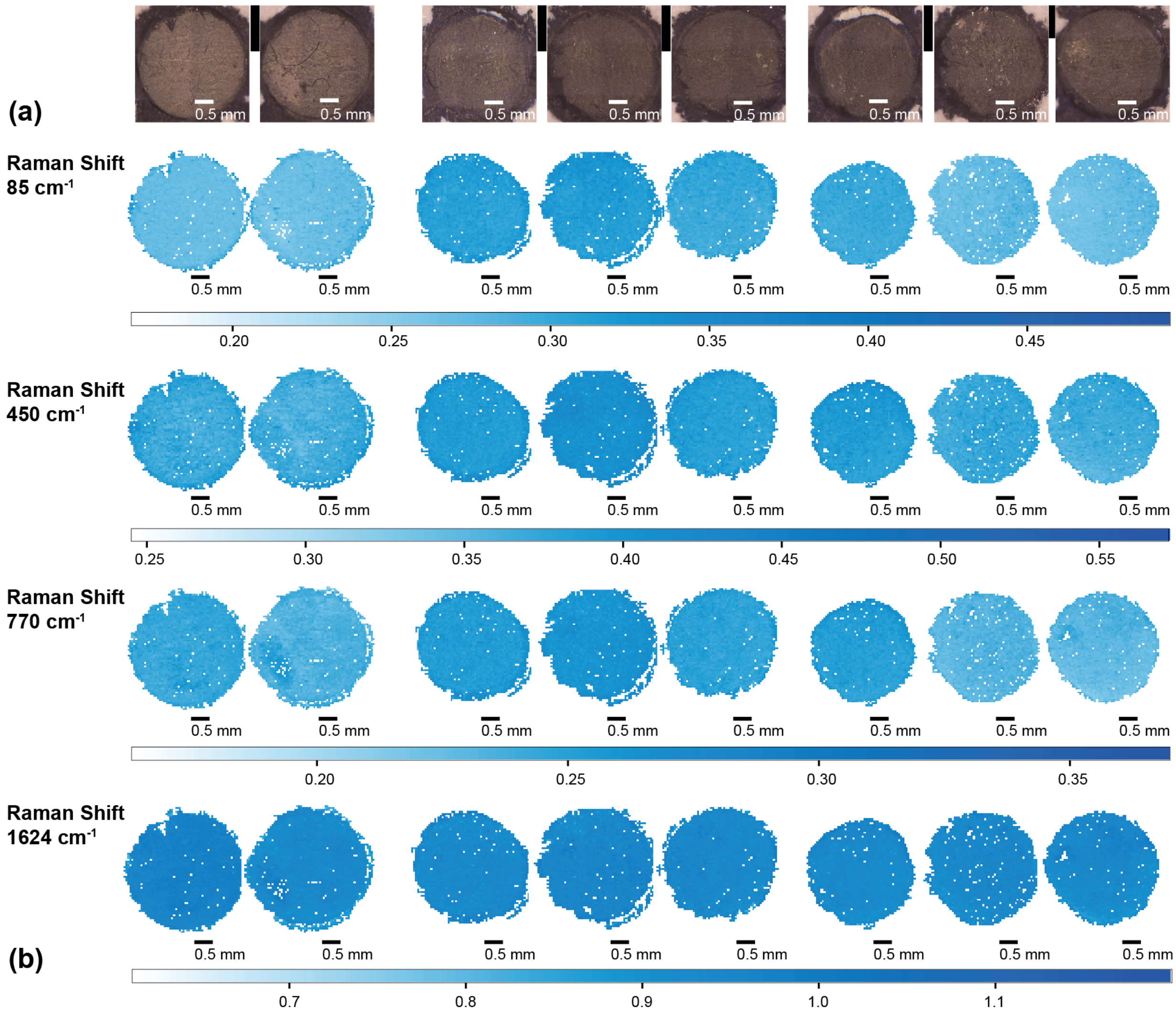


| % RSD | Chip 2 | Chip 3 | Chip 4 | Chip 5 | Chip 6 | Chip 7 | Chip 8 | Chip 9 |
|---|---|---|---|---|---|---|---|---|
| 85 $cm^{-1}$ | 4.05% | 4.27% | 3.22% | 3.15% | 3.41% | 2.32% | 3.22% | 3.11% |
| 450 $cm^{-1}$ | 3.76% | 3.80% | 2.56% | 2.60% | 2.70% | 2.71% | 2.48% | 2.86% |
| 770 $cm^{-1}$ | 2.27% | 3.15% | 2.29% | 2.14% | 2.26% | 2.25% | 2.21% | 2.86% |
| 1624 $cm^{-1}$ | 2.22% | 2.62% | 2.06% | 2.16% | 2.00% | 1.96% | 1.89% | 2.41% |

(c)

| % RSD | Batch 1 | Batch 2 | Batch 3 | Overall |
|---|---|---|---|---|
| 85 $cm^{-1}$ | 10.52% | 4.04% | 5.42% | 8.45% |
| 450 $cm^{-1}$ | 5.72% | 3.52% | 3.62% | 4.91% |
| 770 $cm^{-1}$ | 7.21% | 3.10% | 4.83% | 6.18% |
| 1624 $cm^{-1}$ | 3.03% | 2.18% | 2.12% | 2.54% |

**Supplementary figure 5**. Reproducibility analysis across chips and batches. (a) Optical micrographs of representative silver dendrite sensor chips used in the study. (b) High-resolution Raman mapping of MB at the 85 $cm^{-1}$ quasi-Rayleigh peak, grouped by batches (Batch 1:

Chips 2–3, Batch 2: Chips 4–6, Batch 3: Chips 7–9). This peak reflects the substrate's enhancement effect independent of analyte distribution. (c) RSD% of Raman intensities at 85, 450, 770, and 1624 cm⁻¹ for each chip, each batch, and the entire dataset, demonstrating consistent reproducibility within and across batches.

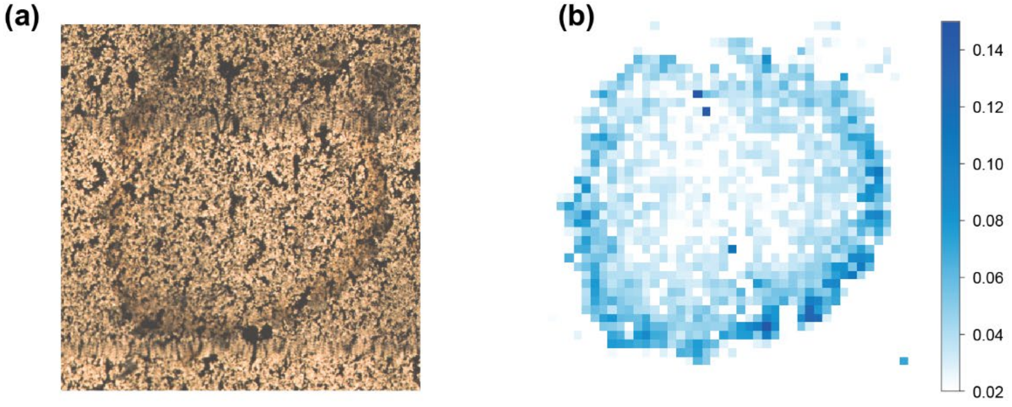


**Supplementary figure 6:** Coffee ring effects in standard silver dendrite substrates. (a) Stitched optical microscope image of a dried 20 µL, 100 nM MB sample on a silver dendrite substrate deposited over a gold-coated glass slide. (b) Raman intensity map at 1624 cm⁻¹ showing higher signal at the droplet edges due to coffee-ring accumulation.

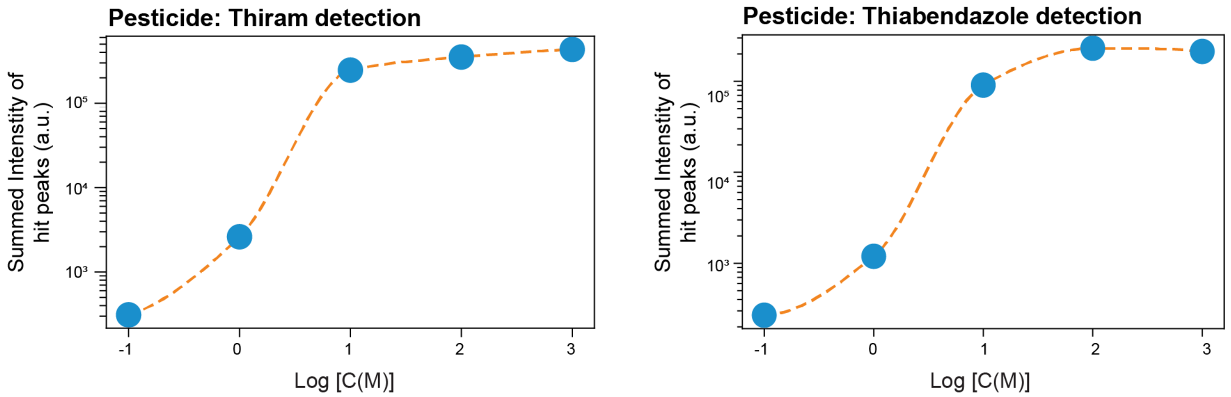


**Supplementary figure 7:** Quantitative SERS analysis of thiram (left) and thiabendazole (right). Summed peak intensity at detected "hit" locations, corresponding to the characteristic peaks at 1383 cm⁻¹ (thiram) and 785 cm⁻¹ (thiabendazole), plotted against concentration.

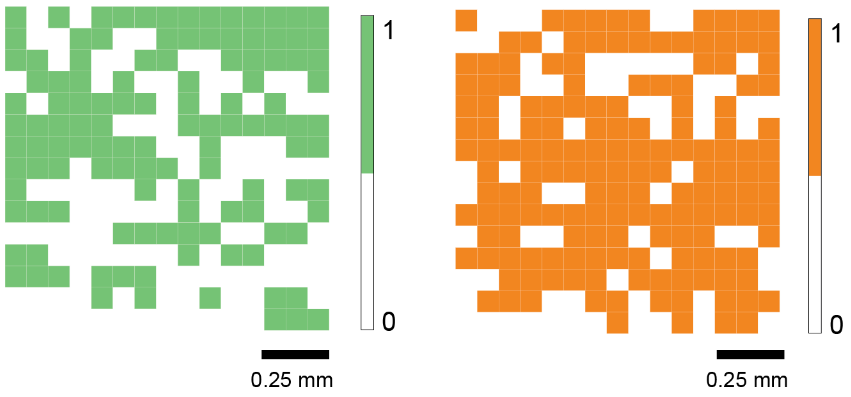


**Supplementary figure 8:** Individual digitized intensity maps for thiram (left) and thiabendazole (right). Maps show spatial "hit" regions for each pesticide, highlighting detection areas identified by the digital SERS method.
